# Simulation of sugar kelp (*Saccharina latissima*) breeding guided by practices to prioritize accelerated research gains

**DOI:** 10.1101/2021.01.21.427651

**Authors:** Mao Huang, Kelly R Robbins, Yaoguang Li, Schery Umanzor, Michael Marty-Rivera, David Bailey, Charles Yarish, Scott Lindell, Jean-Luc Jannink

**Affiliations:** Section on Plant Breeding and Genetics, School of Integrative Plant Sciences, Cornell University, Ithaca, NY, USA; Department of Ecology & Evolutionary Biology, University of Connecticut, Stamford CT, USA; College of Fisheries and Ocean Sciences, University of Alaska Fairbanks, Juneau, Alaska; Applied Ocean Physics and Engineering, Woods Hole Oceanographic Institution, Woods Hole, MA, USA; United States Department of Agriculture - Agriculture Research Service, Ithaca, NY, USA

**Keywords:** sugar kelp, Saccharina latissima, simulation, breeding, genetic gain, genomic selection

## Abstract

The domestication process of sugar kelp in the Northeast U.S. was initiated by selective breeding two years ago. In this study, we will demonstrate how obstacles for accelerated genetic gain can be assessed using simulation approaches that inform resource allocation decisions in our research. Thus far, we have used 140 wild sporophytes (SPs) that were sampled from the northern Gulf of Maine (GOM) to southern New England (SNE). From these SPs, we sampled gametophytes (GPs) and made and evaluated over 600 progeny SPs from crosses among the GPs. The biphasic life cycle of kelp gives a great advantage in selective breeding as we can potentially select both on the SPs and GPs. However, several obstacles exist, such as the amount of time it takes to complete a breeding cycle, the number of GPs that can be maintained in the lab, and whether positive selection can be conducted on farm tested SPs. Using the GOM population characteristics for heritability and effective population size, we simulated a founder population of 1000 individuals and evaluated the impact of overcoming these obstacles on genetic gain. Our results showed that key factors to improve current genetic gain rely mainly on our ability to induce reproduction of the best farm-tested SPs, and to accelerate the clonal vegetative growth of released GPs so that enough GP biomass is ready for making crosses by the next growing season. Overcoming these challenges could improve rates of genetic gain more than two-fold. Future research should focus on conditions favorable for inducing spring and early summer reproduction, and increasing the amount of GP tissue available in time to make fall crosses.

## Introduction

Wild kelp forests in the ocean provide important habitat and ecosystem services. They have also been an important source of human food. Due to climate change and other anthropogenic factors, global kelp populations have faced a drastic decline (Moy and Christie 2012, Wernberg et al. 2019, Bricknell et al. 2020). Now kelp farming is largely replacing wild harvests: over 30 million metric tons of seaweed were harvested in 2018, of which 97% came from farms (FAO, 2020). The import of seaweed raw materials to the U.S. in 2016 was more than 10,000 metric tons (over $73 million, National Marine Fisheries Service Office of Science and Technology 2016; Piconi et al. 2020). Uses include human food, animal feed supplements, and pharmaceutical and cosmetic products (Kim et al. 2015; Kim et al. 2017; Kim et al. 2019; Marine Biotech 2015; Schiener et al. 2015; Wang et al. 2020; Yarish et al. 2017). Growing kelp biomass in the ocean offers a unique opportunity to avoid many of the challenges associated with terrestrial agriculture systems, particularly the growing competition for arable land and freshwater resources. In order to meet the demand of our growing population by 2050, we must use the oceans responsibly to build a thriving seaweed farming industry for the production of carbon-neutral fuels, biochemicals, animal feed, and food (Capron et al. 2020; Kurt et al. 2020).

Kelp cultivation has been established for over 60 years in Asian countries. Most recently, there is growing interest in macroalgal cultivation in Europe, South America, and North America (Buschmann et al. 2017; Grebe et al. 2019; Kim et al. 2019; Geocke et al. 2020). Specifically, there are efforts to selectively breed kelp for large-scale food and bioenergy production (Bjerregaard et al. 2016; Hwang et al. 2019; Valero et al. 2017; Geocke et al. 2020) as well as increased demand for germplasm banking to support future cultivation (Barrento et al. 2016; Wade et al. 2020). The U.S. Department of Energy Advanced Research Projects Agency-Energy (ARPA-E) initiated the Macroalgae Research Inspiring Novel Energy Resources (MARINER) program in order to develop new cultivation, management, and breeding technologies that enable cost-efficient seaweed farming in the large U.S. Exclusive Economic Zone and grow into a global leader in the production of seaweeds. The domestication and breeding of sugar kelp, however, is just beginning.

Kelp has a bi-phasic life cycle (Redmond et al. 2014), which provides unique opportunities for selective breeding since breeders could potentially exert selection pressure on both phases within a single growing cycle (Peteiro et al. 2016, Wade et al. 2020). Genetic markers have been used in crop breeding for some time, primarily exploiting large marker-trait associations (Bernardo 2016). In the last decade, genomic selection (GS) has been adapted by numerous breeding programs due to its ability in predicting breeding values that are immediately used for making selections (Meuwissen et al. 2001, Jannink et al. 2010). The use of genomic selection in terrestrial agriculture and aquaculture breeding has a track record of improving gains by ~10% per generation (Gjedrem et al. 2012). Genomic selection uses a training population with both phenotypic and genotypic information to build a model, which then can be used to predict the genomic estimated breeding value (GEBV) of individuals that are related to the training population. As the development of genetic markers and genotyping individuals becomes less costly compared to phenotyping, GS allows breeders to make selections more efficiently (Heffner et al. 2010).

In 2018, a kelp breeding program was initiated by collecting sporophytes (SPs) from the Gulf of Maine (GOM) to southern New England (SNE). Our primary breeding goal is to improve biomass-related traits including wet weight and percentage dry weight, and to reduce biomass ash content. From the wild-sampled SPs, over 700 uniclonal gametophytes (GPs) were isolated and over 200 of these were grown to sufficient biomass for genotyping and for crossing to create progeny SPs, which were planted and evaluated on nearshore kelp farms. Within a cross, each SP has the exact same genotype, resulting in genetically uniform one-meter line “plots” in the farm. A detailed description is reported in Umanzor et al (2020). In the spring of 2019, the farm-grown SPs were measured, and samples were collected to culture in the lab and induce GPs for the next crossing, planting and harvesting cycle. In our current scheme, we use GS to predict the breeding value of gametophytes, select the best ones, and prioritize crossing these GPs to create new sporophytes. Kelp’s biphasic life cycle (Fig. 1a) allows us to potentially exert tremendous selection pressure on GPs, as we aim to predict combining abilities of parental GPs using the SP performance. This will empower us to prioritize crosses and evaluate SPs that are more likely to become high-performing varieties.

**Figure 1.**
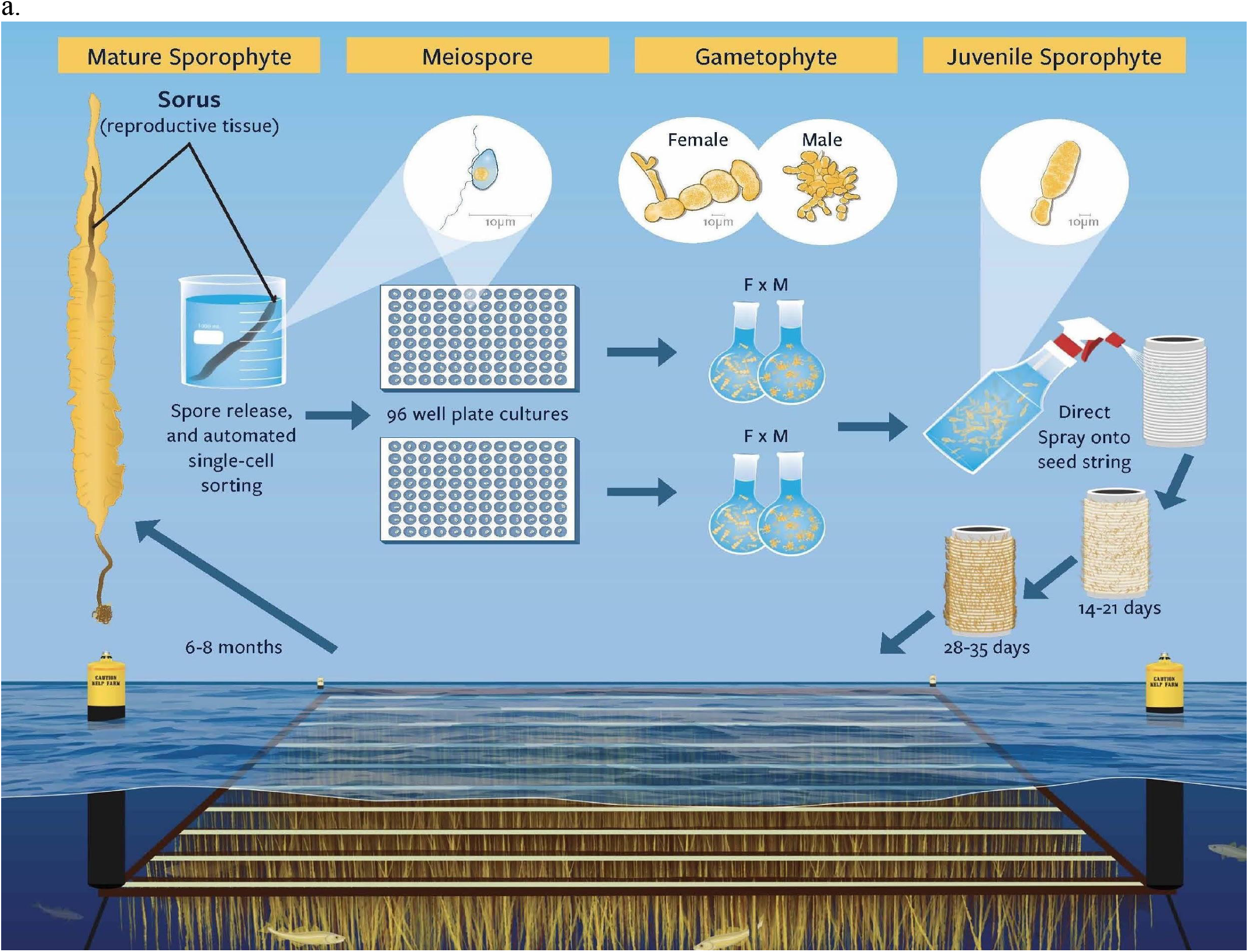
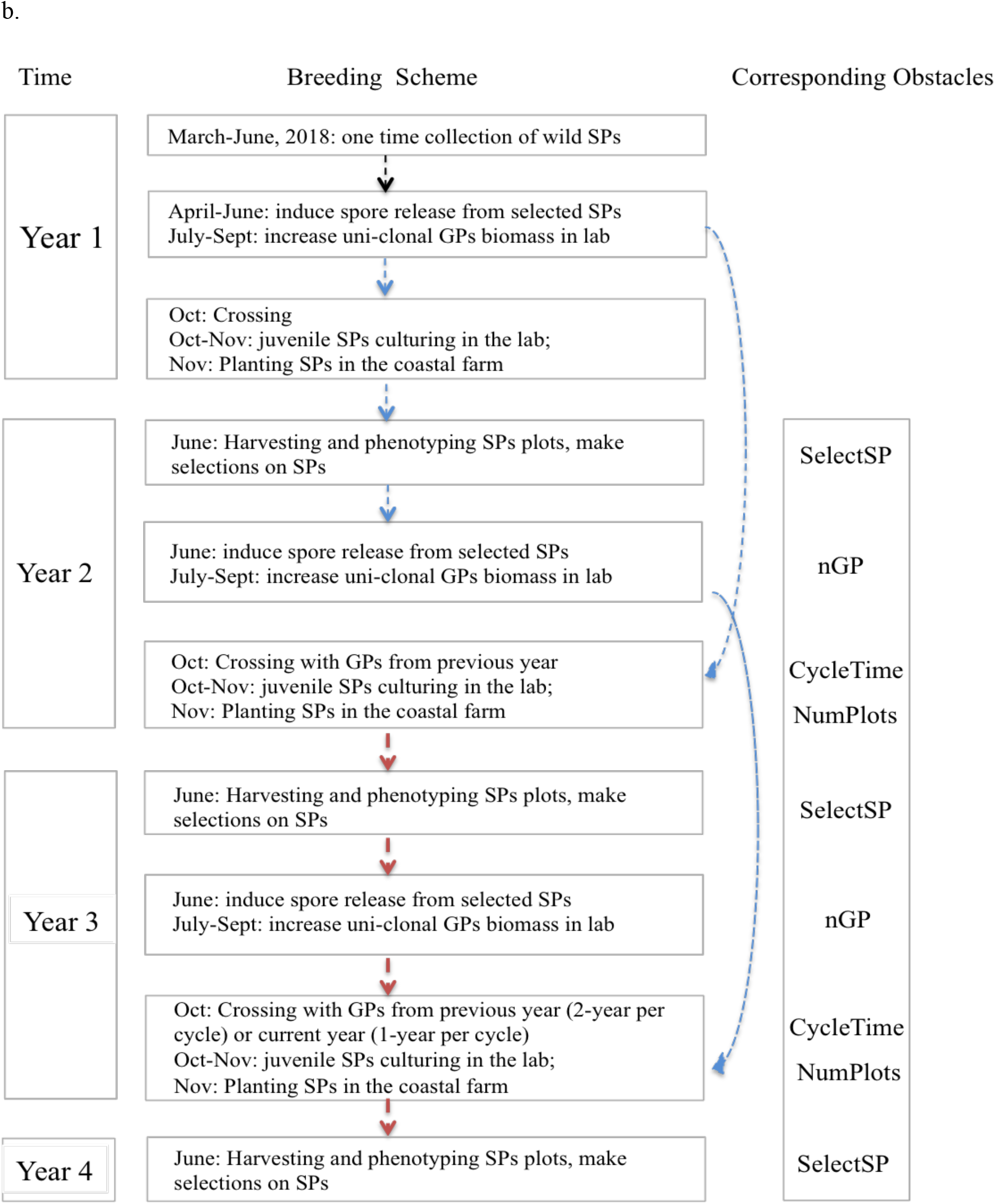
(a.) Biphasic life cycle and breeding pipeline of sugar kelp (*S. latissima*) in our research project. Represented are meiospore release, flow cell sorting to 96-well plates, propagation to sufficient biomass for crossing, spraying of crossed SPs onto seed string, and outplanting to a farm-like common garden field experiment. (b.) Breeding scheme timeline view and the corresponding obstacles on number of GPs (nGP), Number of SP plots evaluated on farm (NumPlots), the selection on SPs (phenotypic vs random selection), and CycleTime (1-year vs 2-year).

Given our experience, we now have a better understanding of significant obstacles to our breeding effort and the investments that might be exerted to overcome those obstacles. To guide the research effort objectively, the extent of accelerated gain from different possible investments and interventions needs to be assessed via simulation. These simulations will help early kelp breeding efforts utilize limited research and development investment for maximal breeding efficiency and genetic gain (the improvement in population genetic mean).

We have identified four obstacles. First, we can collect sorus tissue in the spring from farm-grown SPs and express meiospores that can be individually isolated to grow out to become clonal GPs. However, thus far, we have not routinely succeeded in producing enough clonal GP biomass to make crosses that can generate hundreds of SPs by the fall of the same year. Instead of completing a breeding cycle in one year, our breeding program started with a two year cycle. This slow growth of the clonal GPs represents ***Obstacle 1***. The technical advancement to overcome *Obstacle 1* and complete a breeding cycle in one year would entail some combination of the following:

1. Methods to enhance the growth rate of the GPs so that GPs sampled in the spring have sufficient biomass to make crosses in the fall; or
2. Methods to make crosses that require less GP biomass but that nevertheless produce plots with adequate numbers of SPs.

Currently, we are limited to making no more than 400 crosses per year, due to the labor intensity of maintaining and growing GP cultures in the lab. This bottleneck limits the number of crosses that can be planted and evaluated and represents ***Obstacle 2***. Limiting the number of crosses and associated phenotypic variance can reduce the expected selection intensity and genetic gain. Overcoming *Obstacle 2* would require the ability to maintain and culture more GPs in the lab.

Though we can successfully phenotype and rank SPs after a growing season, we have minimal ability to exert positive selection on them since many of the top ranked SPs did not become reproductive prior to harvest. Consequently, our selection of SPs as parents for the next generation of GPs is limited. The lack of selection pressure applicable to SPs represents ***Obstacle 3***. The biphasic nature of kelp should enable two selection events per breeding cycle, one event on the SPs and one on the GPs they produce. In the absence of selection on the SPs, we currently miss an opportunity for genetic gain. Overcoming *Obstacle 3* would entail rapid identification of top SPs, and artificial laboratory induction of SPs to enter reproductive phase (Pang and Luning, 2004).

Finally, we have shown that it is possible to automate isolating meiospores individually into 96-well plates using flow cytometry (Augyte et al. 2020). This sorting method showed a maximum effectiveness of 76% in gametophyte development (Augyte et al. 2020). We considered the average value in gametophyte survival (i.e., 24 GPs per plate) as a reference parameter in our breeding program. Low GP survival during flow cytometry represents ***Obstacle 4***. Investment in the flow cytometry method to either increase GP survival or enable the preparation of more plates, thus generating more GPs from which to select, would overcome this fourth obstacle.

Using simulation, we aim to compare genetic gain after 5 cycles, examining the impacts of overcoming the aforementioned obstacles. This study will guide our decision-making to optimize resource allocation in the next phase of research, and allow other kelp breeders to focus on advancing these areas most needed.

Simulation studies have been a useful tool in assisting breeders’ decision-making. They are often used to dissect problems that are difficult (expensive or time consuming) to be addressed experimentally. Simulation models can be used to refine more useful experiments based on prior results and experience. For instance, in order to evaluate different ways of improving nitrogen use efficiency for wheat, Dresbøll and Thorup-Kristensen (2014) simulated models mimicking both above and underground plant and environment interactions as well as effects of crop management strategies. These models provided useful guidelines for crop management and variety selection. Simulation results help optimize breeding resource allocation as researchers compare different strategies and predict the potential effects caused by different variables (Parry et al. 2020, Sun et al. 2011, Yamamoto et al. 2016). Simulation approaches were also used to identify the best field experimental design in order to most effectively control for spatial variation in agriculture and forestry studies (Gezan et al. 2010). The selection advantages of GS versus using phenotypic selection were evaluated using simulation approaches for barley (based on real marker data, Iwata and Jannink, 2011) and for *Cryptomeria japonica* (purely simulated data, Iwata et al. 2011). Hickey et al. (2014) simulated breeding schemes incorporating GS and assessed GS accuracies to strategize resources allocated between genotyping versus phenotyping, and between the sizes of populations versus numbers of replications (Lorenz, 2013). The potential genetic gains for a small young sorghum breeding program were assessed via simulation (Muleta et al. 2019).

In aquaculture, breeding simulation studies have also been applied to address a variety of questions (Zenger et al. 2019), including assessing the changes of inbreeding rates over time (Bentsen and Olesen 2002), evaluating the effects of mating strategies on the changes of genetic gain in 10 generations of aquaculture selection (Sonesson and Ødegård, 2016), and assessing the genomic prediction accuracy using either identical by state or identical by descent genomic relationship matrices (Vela-Avitúa et al. 2015). Zenger et al. (2019) reported that at least 36 simulation studies were relevant in aquaculture breeding evaluating different mating designs, selection strategy, family and genome sizes, and their effects on changes of breeding program over different generations.

For simulation studies to be valuable guides, they must be appropriately parameterized. In our breeding work, we have measured various traits at plot and individual levels (Umanzor et al. 2020). Heritability using data across two growing seasons varied among traits and ranged from 0.05 to 0.58, where dry weight per meter and ash free dry weight heritabilities were approximately 0.4. The percent dry weight had the lowest heritability of 0.05. Furthermore, population genetic analyses on the wild samples were performed to understand their diversity, the relationships among them, and their population history in terms of effective population size (Mao et al. 2020). Our simulation parameters were chosen on the basis of these heritability values and on effective population size estimated using founder markers linkage disequilibrium (LD). In this paper, we present a simulation exercise based on these parameters to prioritize research to overcome the obstacles limiting optimum gain from selection.

## Materials and Methods

### Defining the four major obstacles

Sampling the kelp sporophytes in the wild and culturing the founder gametophytes (GPs) was a one-time event and is not counted in the breeding cycle. We define a breeding cycle for sugar kelp starting in the fall of the year when we cross GPs, and ending just before we cross GPs for the following year (Fig. 1b).

*Obstacle 1* is related to the challenge of cultivating enough biomass from GPs collected from farm-evaluated sporophytes (SPs) to make new crosses within the same breeding cycle. In the simulation, we assumed we could reduce the cycle time from two years to one year. *Obstacle 2* is related to limited capacity to grow GPs for crossing. Simulation scenarios assumed we could design space and labor-saving machines for the lab/hatchery phase and manage higher throughput phenotyping to evaluate 1000 plots instead of 400 plots each year. *Obstacle 3* is based on the fact that we were not able to exert selection on farm grown SPs using their phenotypic data because they were not reproductive and we could not harvest their spores and produce the next generation of GPs. In our simulation, we assume the top-ranked sporophytes could be artificially manipulated to be reproductive, hence we could perform phenotypic selection on these sporophytes rather than applying random selection (Pang and Lüning, 2004). Finally, *Obstacle 4* simply affects how many GPs we can collect per parental SP, with our current maximum of 24 but a possible maximum of 96.

### Simulation parameters

#### 1. Founder population characteristics

We first needed an estimate of the effective population size of kelp founders. To obtain this estimate, marker data on 140 wild SPs samples from GOM was generated via DArT technology (Mao et al. 2020). Data cleaning was similar to Mao et al. (2020). Markers were filtered by removing ones with more than 10% missing data and those severely departing from Hardy-Weinberg Equilibrium (P-value < 0.01) in more than 25% of the collection sites. Markers with minor allele frequency less than 0.05 and individuals with more than 50% missing data were also removed. A final set of 4906 markers were retained and imputed using the rrBLUP package A.mat function (Endelman et al. 2011) in R (R Development Core Team, 2018). Linkage disequilibrium between markers was estimated using the genetics package (Warnes et al, 2012). Average LD score were estimated to be 0.08, which were then used in estimating the effective sample size (*N_e_*), according to Sved (1971):

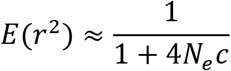

where *E*(*r*^2^) is the expected *r*^2^ for which we used the average LD score of 0.08, and c is the recombination rate among all sites assumed to be 0.5, given that the vast majority of pairs of sites are on different chromosomes. This gives an estimated *N_e_* = 60. We know that the GOM population is strongly structured (Mao et al. 2020), which may cause *N_e_* to be underestimated. Thus we also ran simulations with a setting of *N_e_* = 600. A total of 1000 SP individuals were simulated as our founders with the effective population size of either *N_e_* = 60 or *N_e_* = 600.

The ploidy level was set to 2 and the number of chromosomes was assumed to be 31 based on its close congener *Saccharina japonica* (Liu et al. 2012). Per chromosome, the number of segregating sites and the number of QTL were set to 500 and 100, respectively. These values assume that the trait is polygenic but are otherwise somewhat arbitrary and chosen referring to those in Muleta et al. (2019).

A mixed model, including genetic effects of SPs as random effects, then growth line, blocks, date of harvesting, and reference checks as the fixed effects, was conducted to estimate the narrow sense heritability using 2018-2019 and 2019-2020 two field season GOM farm SP data. Heritability was estimated using:

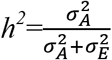

where 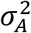 is the estimated additive variance for the SPs and 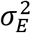 is error variance from a mixed model. Trait heritabilities ranged from 0.05 to 0.50 for plot-level traits and 0.06 to 0.58 for individual-level traits using both years’ data. In the simulation study, the trait genotypic variance was set at 1 and error variance at 4 or 1 so that initial heritability was 0.20 or 0.50 for biomass related traits.

#### 2. Breeding Scheme

Historically, we have been able to produce enough GP biomass to make 2 crosses per GP. Consequently, we assumed that same capacity in the simulation scheme. We created initial SP founder populations of 1000 individuals and allowed each SP to generate two GPs, giving enough GPs to make either 400 or 1000 crosses for downstream generations without exerting selection pressure on the founder population. The simulation program randomly assigned “F” and “M” sexes to GPs generated from the founder population. Ten percent of SPs were selected either randomly or based on phenotype to be parents of the next generation GPs. Thus, 40 and 100 SPs were selected from 400 and 1000 SPs evaluated, respectively. From these selected SPs, we assumed flow cytometry would be used to obtain GPs from each SP (Augyte et al. 2020). This automated spore sorting technology produces viable uni-clonal isolations on average in 25% of the wells of a 96-well plate (i.e., 24 GPs) from spores released from an individual SP. An ideal situation where all 96 GPs in the plate are viable was also included in the simulation scheme. Generating 96 GPs means there will be four times more GPs to select from to make crosses for SPs, enabling higher selection intensity. The farm-evaluated SPs phenotypic data from both years was used to train a genomic selection model which was used to predict the breeding values of all GPs coming out of the flow cytometry process. Either 200 or 500 top ranked GPs based on their predicted breeding values would be selected to make the 400 or 1000 crosses as farm-evaluated SPs plots. Note that with this scheme, changing the number of SPs evaluated does not change the selection intensity either during SP or GP selection, whereas changing the number of GPs generated changes the selection intensity during the GP selection stage.

#### 3. Estimating Genetic gain over 5 breeding cycles

Breeding scheme simulation was done using the AlphaSimR package in R (Faux et al. 2016). Each scheme was simulated 100 times, and the average genetic gain as well as genetic variance at each GP stage was calculated over 5 cycles of selection. Because we were mainly interested in evaluating the trend of genetic gain from different breeding schemes, the reference point for genetic gain could either be for GPs or SPs. We used GPs.

## Results

### Simulation output

The ability to exert selection on the farm-evaluated SPs (SelectSP), the number of years per breeding cycle time (CycleTime), and the number of gametophytes per SP surviving the flow cell cytometry system (nGP) were the three significant contributors to the changes of genetic mean over time (Table1). We did not observe significant interactions between these factors (Table 1).

**Table 1.** ANOVA on genetic mean split by founder effective population size (*N_e_*) and heritability (*h^2^*).

). a. $N_e = 60$ , $h^2 = 0.5$
|  | Df | Sum Sq | Mean Sq | *F* | *P-value* |
| --- | --- | --- | --- | --- | --- |
| SelectSP† | 1 | 47.6 | 47.6 | 22.4 | 7.34E-06*** |
| NumPlots | 1 | 1.0 | 1.0 | 0.5 | 0.496 |
| CycleTime | 1 | 13.7 | 13.7 | 6.4 | 0.013* |
| nGP | 1 | 10.0 | 10.0 | 4.7 | 0.032* |
| SelectSP:NumPlots | 1 | 0.0 | 0.0 | 0.0 | 0.927 |
| SelectSP:CycleTime | 1 | 0.5 | 0.5 | 0.3 | 0.613 |
| SelectSP:nGP | 1 | 0.4 | 0.4 | 0.2 | 0.648 |
| NumPlots:CycleTime | 1 | 0.1 | 0.1 | 0.0 | 0.876 |
| NumPlots:nGP | 1 | 0.1 | 0.1 | 0.0 | 0.830 |
| CycleTime:nGP | 1 | 0.4 | 0.4 | 0.2 | 0.663 |
| Residuals | 101 | 215.0 | 2.1 |  |  |

b. $N_e = 600$ , $h^2 = 0.5$
|  | Df | Sum Sq | Mean Sq | *F* | *P-value* |
| --- | --- | --- | --- | --- | --- |
| SelectSP† | 1 | 48.5 | 48.5 | 31.1 | 2.02E-07*** |
| NumPlots | 1 | 0.4 | 0.4 | 0.3 | 0.614 |
| CycleTime | 1 | 10.8 | 10.8 | 6.9 | 0.010** |
| nGP | 1 | 6.9 | 6.9 | 4.4 | 0.038* |
| SelectSP:NumPlots | 1 | 0.0 | 0.0 | 0.0 | 0.897 |
| SelectSP:CycleTime | 1 | 0.4 | 0.4 | 0.3 | 0.610 |
| SelectSP:nGP | 1 | 0.4 | 0.4 | 0.3 | 0.614 |
| NumPlots:CycleTime | 1 | 0.0 | 0.0 | 0.0 | 0.902 |
| NumPlots:nGP | 1 | 0.1 | 0.1 | 0.0 | 0.838 |
| CycleTime:nGP | 1 | 0.3 | 0.3 | 0.2 | 0.666 |
| Residuals | 101 | 157.2 | 1.6 |  |  |

c. $N_e = 60$ , $h^2 = 0.2$
|  | Df | Sum Sq | Mean Sq | *F* | *P-value* |
| --- | --- | --- | --- | --- | --- |
| SelectSP† | 1 | 19.3 | 19.3 | 15.4 | 0.000*** |
| NumPlots | 1 | 1.2 | 1.2 | 0.9 | 0.339 |
| CycleTime | 1 | 8.1 | 8.1 | 6.5 | 0.012* |
| nGP | 1 | 6.7 | 6.7 | 5.3 | 0.023* |
| SelectSP:NumPlots | 1 | 0.0 | 0.0 | 0.0 | 0.967 |
| SelectSP:CycleTime | 1 | 0.2 | 0.2 | 0.1 | 0.726 |
| SelectSP:nGP | 1 | 0.2 | 0.2 | 0.2 | 0.660 |
| NumPlots:CycleTime | 1 | 0.0 | 0.0 | 0.0 | 0.849 |
| NumPlots:nGP | 1 | 0.1 | 0.1 | 0.1 | 0.733 |
| CycleTime:nGP | 1 | 0.2 | 0.2 | 0.2 | 0.676 |
| Residuals | 101 | 126.6 | 1.3 |  |  |

d. $N_e = 600$ , $h^2 = 0.2$
|  | Df | Sum Sq | Mean Sq | *F* | *P-value* |
| --- | --- | --- | --- | --- | --- |
| SelectSP<sup>†</sup> | 1 | 19.4 | 19.4 | 21.4 | 1.11E-05*** |
| NumPlots | 1 | 0.4 | 0.4 | 0.5 | 0.486 |
| CycleTime | 1 | 6.8 | 6.8 | 7.6 | 0.007** |
| nGP | 1 | 4.3 | 4.3 | 4.7 | 0.032* |
| SelectSP:NumPlots | 1 | 0.0 | 0.0 | 0.0 | 0.897 |
| SelectSP:CycleTime | 1 | 0.3 | 0.3 | 0.3 | 0.599 |
| SelectSP:nGP | 1 | 0.1 | 0.1 | 0.2 | 0.691 |
| NumPlots:CycleTime | 1 | 0.0 | 0.0 | 0.0 | 0.827 |
| NumPlots:nGP | 1 | 0.1 | 0.1 | 0.1 | 0.769 |
| CycleTime:nGP | 1 | 0.1 | 0.1 | 0.1 | 0.726 |
| Residuals | 101 | 91.4 | 0.9 |  |  |

\* $P < 0.05$ , \*\* $P < 0.001$ , \*\*\* $P < 0.0001$
<sup>†</sup> SelectSP: Selection among SP based on phenotype or at random. NumPlots: Common garden of 400 versus 1000 field plots. CycleTime: 1-year versus 2-year cycle. nGP: number of GPs obtained per parental SP of 24 or 96.

The baseline simulation scheme represented our current state of the art, where 400 SPs are phenotyped in the field, no selection pressure is exerted on the SPs, and the breeding cycle takes 2 years. Our number of GPs per SP was either 24 or 96. The changes from Fig. 2a to Fig. 2b reflect the effects of overcoming *Obstacle 4* where higher nGP could be obtained through a single cell sorting flow cytometry step (nGP=24 versus nGP=96). This change from nGP=24 to nGP=96 led to a gain increase of 37% averaged across all other factors (nGP, Table 1, Fig. 2). Relative to the baseline, the ability to exert selection on SPs (*Obstacle 3*) and decreasing the breeding cycle time (*Obstacle 1*) led to gain increases of 101% and 45%, respectively, averaged across all other factors. Though the effect of increasing the number of plots phenotyped was not statistically significant (NumPlots, Table 1), numerically this change increased gain by an average of 11% (overcoming *Obstacle 2*). We did not observe significant interactions: the effects of overcoming each obstacle were additive (Table 1), and overcoming all four obstacles led to the greatest gain (Fig. 2). Heritability also played a role in affecting the genetic gain (Table 1), where *h^2^*=0.5 generated higher genetic mean after 7 years of breeding than *h^2^*=0.2 (Figs. 2a and 2b). This trend was consistent regardless of the number of gametophytes or effective population size.

**Figure 2.**
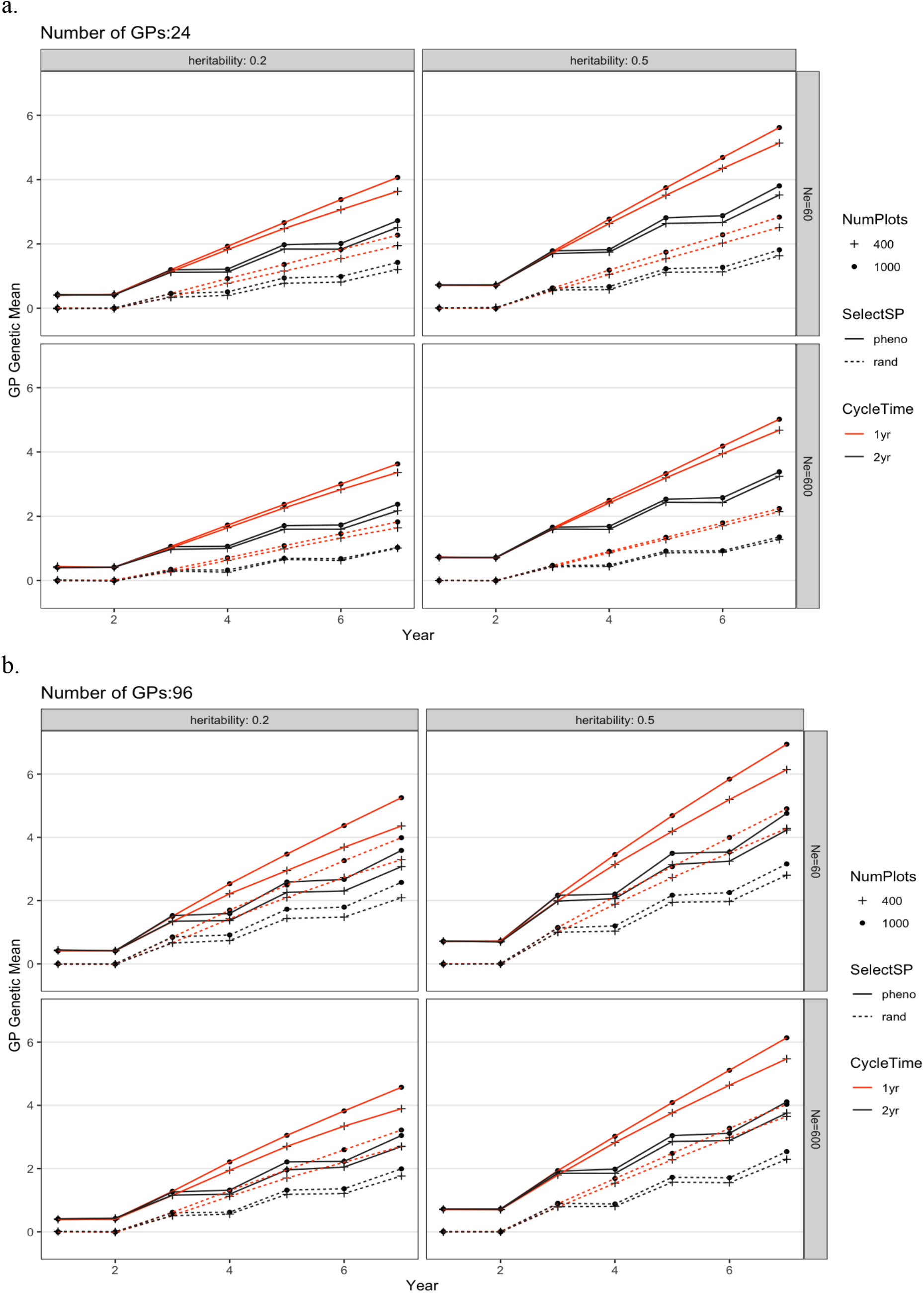
Genetic mean from different breeding schemes over 7 years. The routine breeding scheme starts in year 3. Each figure shows NumPlots: Evaluate 400 versus 1000 plots; SelectSP: phenotypically select the best (pheno) versus random (rand) sporophytes for producing new crosses; and CycleTime: 1-year (1yr) versus 2-year (2yr). Subpanels separate different founder population effective population sizes of 60 (Ne60) and 600 (Ne600) and trait heritabilities of *h^2^* =0.5 and *h^2^* =0.2 when a.) 24 or b.) 96 gametophytes were propagated from each parental SP. Each scheme was repeated 100 times and genetic values shown were averages. The standard error was smaller than the figure symbols and is not shown.

The breeding scheme interventions simulated also affected the genetic variance remaining after seven years of improvement (Fig. 3). All three interventions that significantly increased genetic gain also caused decreases in genetic variance. The smallest change in genetic variance occurred as a result of selecting SPs on phenotype. Selection causes variance decreases both because of the Bulmer effect and because high fitness ancestors contribute disproportionately to descendants. With the 1-year per cycle scheme, the population went through twice as many selection events as with the 2-year scheme, leading to a greater decrease in genetic variation over the seven years (Fig. 3, Online Resource 1). For all combinations of other factors, there was a higher final genetic variance when nGP was 24 than when it was 96. The increased selection intensity from this intervention caused a greater variance decrease than for any other intervention. The only intervention that caused increased final genetic variance was evaluating more SP plots per year (1000 versus 400). In this case, increasing the number of plots caused increased effective population size and thus greater maintenance of variance. It also caused increased genomic prediction accuracy, which has also been shown to maintain genetic variance (Jannink et al. 2010). These effects are depicted in Figure 4. The low level of interaction between simulated factors can also be seen in Figure 4 by the fact that lines linking simulation settings with and without the interventions are approximately parallel and of similar length, indicating that changing one factor has basically the same effect regardless of the levels of the other factors.

**Figure 3.**
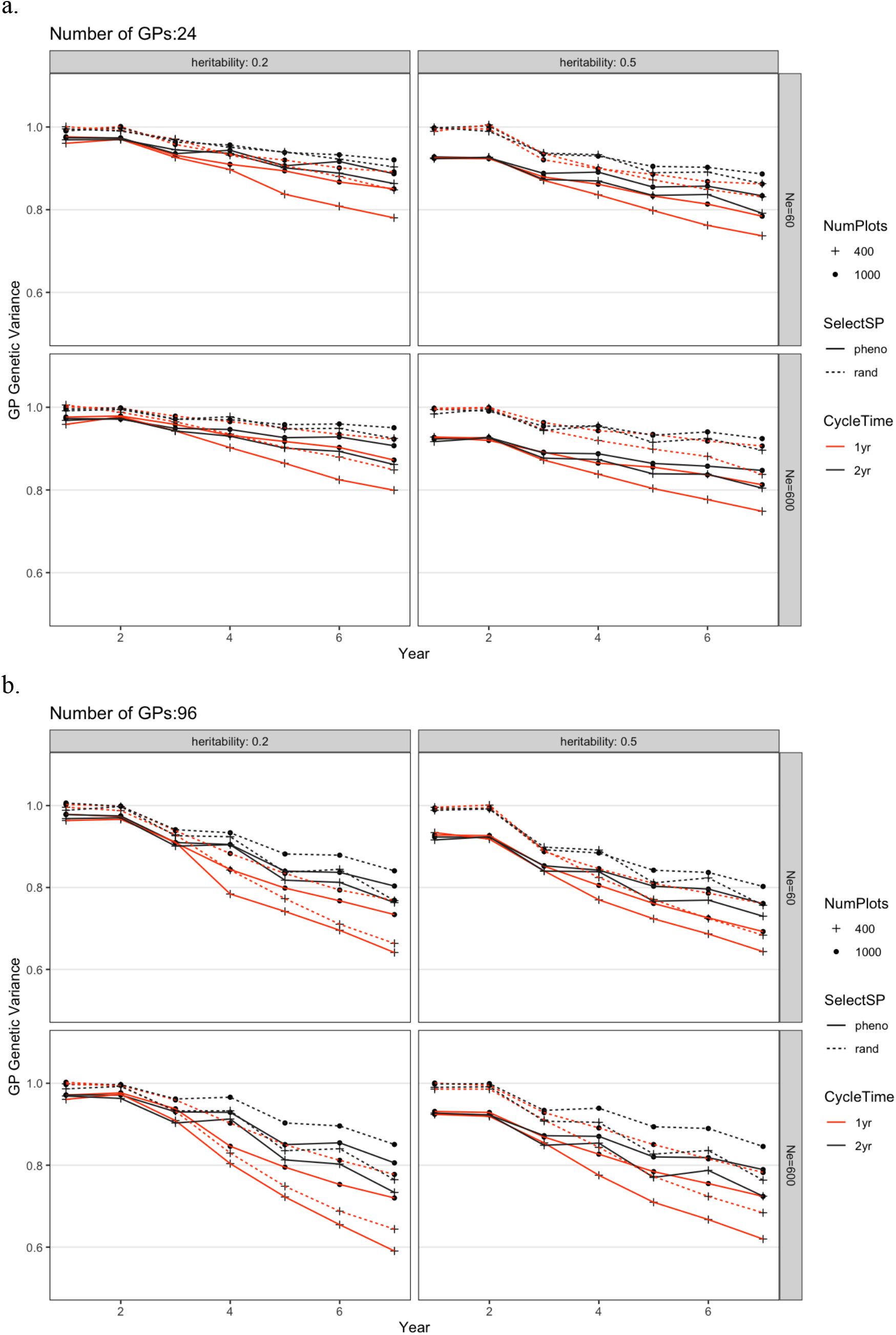
Change of genetic variance from different breeding schemes over 7 years for a.) 24 or b.) 96 gametophytes per parental sporophyte. The scheme abbreviations are the same as for Figure 2. Each scheme was repeated 100 times and genetic variance shown was the average. The standard error was smaller than the figure symbols and is not shown.

**Figure 4.**
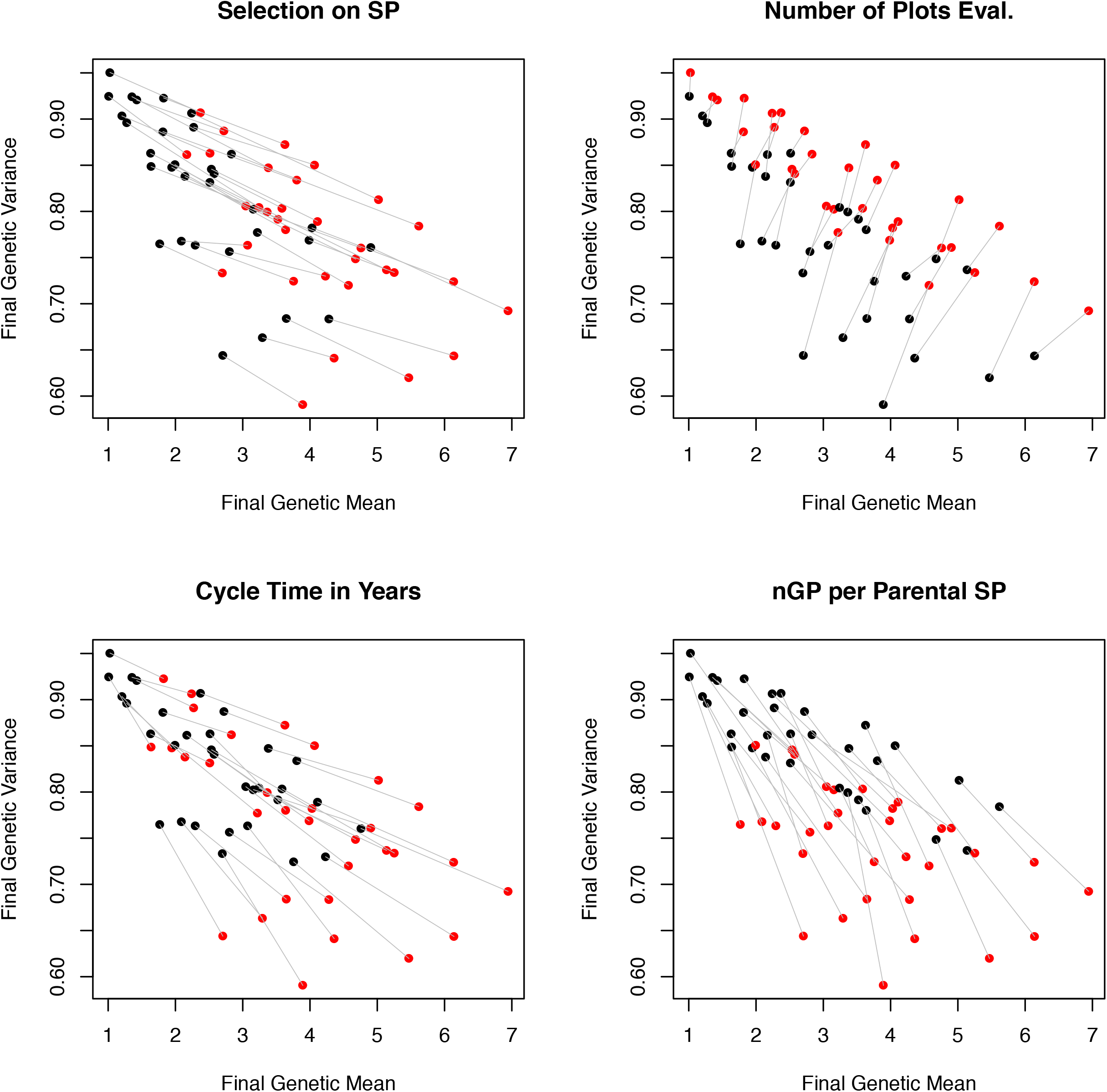
Four views of the simulation results on the final genetic variance and genetic mean. Each view presents the same scatterplot, with each point representing the mean outcome of 100 simulations of one scheme. Each view shows a different obstacle to overcome, with the color of the point determined by the current practice (black) or the improved practice (red). Gray lines connect simulation schemes that are identical except for this change in practice.

## Discussion

### Which obstacle should we focus on?

Simulation is a useful tool for guiding researchers in decision-making especially for young breeding programs (Muleta et al. 2019) and in assisting selection strategy and decision-making processes (Zenger et al. 2019). We simulated different breeding schemes, each overcoming a major obstacle we have encountered in two seasons of kelp breeding. Assessing which scheme generated the highest genetic gain allows us to prioritize research directions and derive the most benefit from a limited budget.

The simulation revealed a robust result that the highest genetic gain can be achieved by exerting selection on SPs phenotypically (overcoming *Obstacle 3*), and then by reducing the time for obtaining sufficient GP biomass such that a one-year cycle is enabled compared to our current two-year cycle (overcoming *Obstacle 1*). Increasing the number of viable GP we obtain per parental SP (overcoming *Obstacle 4*) also delivered significant gain, while, somewhat surprisingly, phenotyping more SP plots (overcoming *Obstacle 2*) did not. These conclusions were not affected by the founder population effective population size or the trait heritability. Thus, the clear direction to prioritize breeding enhancement is to induce SP spore release and to modify GP culture to accelerate growth. In addition, we should experiment with the amount of GP biomass needed to make sufficient SP progeny. We may not need a full one-meter of line we use as an evaluation plot.

Developing the ability to induce top performing SPs to release spores (*Obstacle 3*) can be a challenge for the following reasons. First, typically only approximately 10% of the plots are fertile at the optimum time of harvest as measured by most marketable yield. These crosses are not necessarily the top performing ones. Ideally, we aim to select crosses in the top 10% for performance and artificially induce them indoors if necessary. This has proven successful on a small scale if desirable SPs are identified within a day of harvest. Overcoming *Obstacle 3* requires greater investment in labor to identify and separate candidate SPs, and investment in culture space.

Our second best option is to accelerate GP growth by overcoming *Obstacle 1*, which could also be the hardest task. In brief, it takes four to eight weeks to induce immature SPs to full maturity and release meiospores under artificial conditions in the lab (Pang and Lining 2004, Flavin et al. 2013, Remond et al. 2014). Once meiospores are released, flow-cytometry techniques can be implemented to isolate single-cell gametophytes into 96-well plates. A second isolation is performed approximately two to four months later when GPs develop into tufts large enough (>100 μm), to be sexed and moved to individual Petri dishes for filament fragmentation. Once sufficient uniclonal biomass is achieved (~10 mg to cover 1 m plots), which can take up to another four months, crosses are made by mixing female and male GPs at a 2:1 ratio (Umanzor et al. 2020, Fig. 1). Outplanting at sea occurs 4-6 weeks following SP attachment onto the seed string (Flavin et al. 2013, Remond et al. 2014). Overall, this process of uniclonal GP isolation, growth and crossing is effective but typically requires 12 months, in contrast to the six months between optimal kelp harvesting (end of May to early June) to crossing and outplanting (November to December).

Possible means of accelerating GP growth include optimizing lighting, nutrient and temperature regimes, as well as novel biomass fragmentation protocols. It might be possible to optimize GP biomass development by transferring them earlier to plates with bigger wells (i.e. from 96-well plates to 24-well plates) that would allow better light penetration. Generally, GP growth is limited by the natural biological programming of cell division and a propensity to self-shade in its puff-ball growth form. However, some GPs grow faster than others, and selecting for GP growth performance could be incorporated in the breeding program. In order to test and see if 1-year cycle time is feasible in our current breeding program (approximately six months between GP isolation and crossing), we are experimenting with using a minimum amount of biomass to make crosses and generate at least a single SP blade. The function of this blade would not be for evaluation of SP performance but for recombining the best GPs in the hope of getting improved recombinants. The approach will generate phenotypic data on the individual SP but not on biomass per meter of line, which is a plot-level trait. Hence this procedure would not be a full representation of the simulated 1-year per cycle scheme. Nonetheless, this will be a proof of concept for us to accelerate the GP culturing process.

Another possibility that is used in forage breeding (Resende et al. 2013) would be to evaluate segregating plots, in our case created from crossing multiple female GPs from one SP with multiple male GPs from another SP. The between plots variance for such mixed plots would be less than that for the uniform SPs plots. Furthermore, maintaining multiple individual GPs only until they can be sexed and co-cultured together would reduce some labor. Such mixed plots would generate sufficient biomass more quickly to facilitate one-year breeding cycles.

Overcoming *Obstacle 2* by increasing the number of GPs per parental SP can potentially be done easily. A simple approach would be to increase the number of plates automatically sorted by flow cytometry per parental SP, which would increase the number of GPs in the genomic selection step, allowing higher selection intensity. Nonetheless, this would result in increasing the number of cultures to maintain in the lab, which leads to more labor and cost. The use of flow cytometry sorting expedites the initial isolation process but the parameters determine the survival of spores is not well understood. The condition of sorus tissue prior to spore release and sorting likely has an effect on spore viability. Percentage viability varied across samples presumably because of differences in sorus tissue condition and handling prior to sorting (Augyte et al, 2020). An issue that should be investigated is whether the selection pressure caused by flow cytometry mortality has pleiotropic effects that might negatively affect SP growth or reproduction. If not, the mortality should generate its own natural selection response that will eventually mitigate this obstacle.

Increasing the number of plots (from 400 to 1000) could be accomplished without new research, but could be costly since it would require more GP grow out space and labor. This change generated only a small increase in the rate of genetic gain. An additional benefit to increasing the number of SPs being phenotyped, however, was that it maintained genetic diversity and slowed down the decrease of genetic variance (Fig. 3, Fig. 4). The proportion of GPs selected out of SPs were the same regardless of testing 400 or 1000 plots, hence increasing the number of plots did not change the selection differential. It did, however, affect the training population size of GS models when selecting new generations of GPs. Larger training population size usually contributes to increased GS accuracy (Poland et al. 2012; Huang et al. 2016). In this case, the increased phenotypic data led to an improved genomic prediction model and its ability to distinguish among-family versus within family effects. That ability can decrease the co-selection of relatives leading to greater maintenance of genetic variation (Jannink et al. 2010). Interestingly, every intervention that led to greater genetic gain also led to greater loss of genetic variance for all changes in practice (Selection on SP, Cycle Time, nGP per parental SP), *except* increasing the number of phenotyped plots which had both increased gain and decreased variance lost (Fig. 4). We also observed in some cases that the principal effect of increasing the number of plots was to cause greater variance to be retained, without increasing the gain from selection substantially (in Fig. 4 the gray lines were close to vertical). Hence, it seems likely that this intervention would benefit our breeding program over the long term.

While in this discussion we have treated heritability as fixed, that is not strictly true. Heritability might be increased if we could improve our planting technique to ensure that plots are more uniformly covered by SPs, so that we obtain successful and uniform growth of SPs in the field. Not surprisingly, higher heritability leads to greater final gain (Figs. 2 and 3). The decreasing trend of genetic variance was expected leading to a relationship where higher final genetic gain coincided with lower genetic variance. It is important to maintain the diversity while we improve the progeny performance (Heffner et al. 2009; Lin et al. 2016). Overall, the robustness of these simulation findings should give us confidence in the research directions they suggest. We believe that these priorities will greatly help accelerate genetic gain in breeding programs and therefore increase the value of kelp farming in the United States and globally.

## Declarations

## Acknowledgements

We acknowledge the funding support from the U.S. Dept. of Energy ARPA-E (DE-AR0000915). We thank Natalie Renier from WHOI Communications Department assisting us to make Figure 1.

## Ethics approval

This research complies with the current laws of the United States of America

## Consent to participate

Not applicable

## Consent to publication

All authors read and approved the manuscript for publication

## Availability of data and material

Supplemental Table is available as Online Resource 1.

## Code availability

Simulation codes are constructed in R using package AlphaSimR

## Authors’ contributions

MH performed the analyses, wrote the manuscript draft and revised the manuscript together with other coauthors. KRR and J-LJ guided the analyses. J-LJ contributed to analysis scripts. YL, SU, SL and CY helped edit the manuscript. YL, SU, MMR, CY, DB, and SL collected phenotypic data, from which the simulation parameters in this manuscript were estimated. SL, CY, and J-LJ led the project and all authors read and approved the manuscript.

