## Supplemental Table 1 for "Simulation of sugar kelp (*Saccharina latissima*) breeding guided by practices to prioritize accelerated research gains"

Supplemental Tables

Supplemental Table 1. ANOVA on genetic variance split by founder effective population size ($N_e$) and heritability ($h^2$).

a. $N_e = 60$, $h^2 = 0.5$

| | Df | Sum Sq | Mean Sq | *F* | *P-value* |
|---|---|---|---|---|---|
| SelectSP† | 1 | 0.1 | 0.1 | 16.7 | 0.000 *** |
| NumPlots | 1 | 0.0 | 0.0 | 1.4 | 0.237 |
| CycleTime | 1 | 0.0 | 0.0 | 3.7 | 0.058 |
| nGP | 1 | 0.1 | 0.1 | 11.7 | 0.001*** |
| SelectSP:NumPlots | 1 | 0.0 | 0.0 | 0.1 | 0.790 |
| SelectSP:CycleTime | 1 | 0.0 | 0.0 | 0.0 | 0.854 |
| SelectSP:nGP | 1 | 0.0 | 0.0 | 0.1 | 0.721 |
| NumPlots:CycleTime | 1 | 0.0 | 0.0 | 0.1 | 0.813 |
| NumPlots:nGP | 1 | 0.0 | 0.0 | 0.0 | 0.828 |
| CycleTime:nGP | 1 | 0.0 | 0.0 | 0.2 | 0.697 |
| Residuals | 101 | 0.6 | 0.0 | | |

b. $N_e = 600$, $h^2 = 0.5$

| | Df | Sum Sq | Mean Sq | *F* | *P-value* |
|---|---|---|---|---|---|
| SelectSP† | 1 | 0.1 | 0.1 | 29.0 | 0.000*** |
| NumPlots | 1 | 0.0 | 0.0 | 5.9 | 0.017* |
| CycleTime | 1 | 0.0 | 0.0 | 4.6 | 0.035* |
| nGP | 1 | 0.1 | 0.1 | 13.6 | 0.000*** |
| SelectSP:NumPlots | 1 | 0.0 | 0.0 | 0.0 | 0.965 |
| SelectSP:CycleTime | 1 | 0.0 | 0.0 | 0.0 | 0.875 |
| SelectSP:nGP | 1 | 0.0 | 0.0 | 0.3 | 0.560 |
| NumPlots:CycleTime | 1 | 0.0 | 0.0 | 0.4 | 0.540 |
| NumPlots:nGP | 1 | 0.0 | 0.0 | 0.6 | 0.437 |
| CycleTime:nGP | 1 | 0.0 | 0.0 | 0.9 | 0.356 |
| Residuals | 101 | 0.5 | 0.0 | | |

c. $N_e = 60$, $h^2 = 0.2$

| | Df | Sum Sq | Mean Sq | *F* | *P-value* |
|---|---|---|---|---|---|
| SelectSP† | 1 | 0.0 | 0.0 | 4.6 | 0.034* |
| NumPlots | 1 | 0.0 | 0.0 | 2.7 | 0.102 |
| CycleTime | 1 | 0.0 | 0.0 | 6.3 | 0.013* |
| nGP | 1 | 0.1 | 0.1 | 15.4 | 0.000*** |
| SelectSP:NumPlots | 1 | 0.0 | 0.0 | 0.0 | 0.912 |
| SelectSP:CycleTime | 1 | 0.0 | 0.0 | 0.1 | 0.786 |
| SelectSP:nGP | 1 | 0.0 | 0.0 | 0.0 | 0.887 |
| NumPlots:CycleTime | 1 | 0.0 | 0.0 | 0.4 | 0.521 |

| | | | | | |
|---|---|---|---|---|---|
| NumPlots:nGP | 1 | 0.0 | 0.0 | 0.5 | 0.496 |
| CycleTime:nGP | 1 | 0.0 | 0.0 | 0.7 | 0.413 |
| Residuals | 101 | 0.6 | 0.0 | | |

d. $N_e = 600$, $h^2 = 0.2$

| | Df | Sum Sq | Mean Sq | *F* | *P-value* |
|---|---|---|---|---|---|
| SelectSP† | 1 | 0.0 | 0.0 | 5.5 | 0.021* |
| NumPlots | 1 | 0.0 | 0.0 | 6.1 | 0.015* |
| CycleTime | 1 | 0.0 | 0.0 | 6.4 | 0.013* |
| nGP | 1 | 0.1 | 0.1 | 20.5 | 0.000*** |
| SelectSP:NumPlots | 1 | 0.0 | 0.0 | 0.0 | 0.978 |
| SelectSP:CycleTime | 1 | 0.0 | 0.0 | 0.0 | 0.950 |
| SelectSP:nGP | 1 | 0.0 | 0.0 | 0.0 | 0.990 |
| NumPlots:CycleTime | 1 | 0.0 | 0.0 | 0.7 | 0.418 |
| NumPlots:nGP | 1 | 0.0 | 0.0 | 0.7 | 0.406 |
| CycleTime:nGP | 1 | 0.0 | 0.0 | 1.4 | 0.238 |
| Residuals | 101 | 0.6 | 0.0 | | |

* P<0.05,** P<0.001, *** P<0.0001

† SelectSP: Selection among SP based on phenotype or at random. NumPlots: Common garden of 400 versus 1000 field plots. CycleTime: 1-year versus 2-year cycle. nGP: number of GPs obtained per parental SP of 24 or 96.
